# A neural network with key-value episodic memory retrieves and organizes memories based on causal event structures

**DOI:** 10.1101/2025.09.01.673596

**Authors:** Hayoung Song, Qihong Lu, Tan T. Nguyen, Janice Chen, Yuan Chang Leong, Monica D. Rosenberg, ShiNung Ching, Jeffrey M. Zacks

## Abstract

Humans reflect on memories to make sense of ongoing events. Past work has shown that people retrieve causally related memories during comprehension, but the mechanisms underlying this process remain unclear. Here, we used a recurrent neural network augmented with a key-value episodic memory buffer and trained it to predict upcoming scenes while watching a television episode. At each time step, the model transformed the current scene into a value representing memory content and a key representing memory address, both stored as episodic memory. The model retrieved selective past values by applying self-attention over stored keys and integrated these memories with the current scene representation to generate predictions. The model retrieved memories similar to those retrieved by human participants watching the same episode during fMRI. Importantly, this similarity disappeared when causal relationships between events were controlled for. The model also represented causally related events with similar patterns, similar to how the human brain represents these events. These findings suggest that using two distinct memory representations allows the model to retrieve memories and organize events based on causal relationships, beyond semantic or perceptual similarities. Together, this work proposes a key-value episodic memory system as a candidate computational mechanism for how humans retrieve causally related memories to comprehend naturalistic events.

## Introduction

Understanding ongoing experience is not just about making sense of what is happening now. People reason about why an event has happened by remembering events that occurred in the past. Since the 1970s, theories have described narrative comprehension as a process of constructing a coherent representation of a situation by inferring cause-and-effect relationships between events^1^. Causal inference occurs throughout comprehension, as people retrieve causally related past events to understand and integrate them with the current event^2–4^.

Initial evidence for this view came from work that asked participants to describe their thought process during comprehension or recall story events afterwards. When participants were asked to think aloud while reading a story, they were more likely to mention past events that were causally related to the current event^5^. Studies also found that events that are more causally connected to other events were more likely to be remembered^6–9^. Complementing these behavioral findings, fMRI work showed that neural patterns of related past events were reinstated in the brain during narrative comprehension^10,11^. Recent work specifically showed that *causally related* past events were reinstated in the brain, more so than semantically or perceptually similar events^12^.

These findings suggest that people reason about causal relationships between events through memory retrieval. But what are the computational underpinnings of this causal reasoning process? In other words, how does the human brain search through a vast store of memories to identify and retrieve causally related ones, and then integrate them with an ongoing event to update situational representations? Through this memory retrieval and integration process, how does the brain come to represent events based on their causal relationships? These questions call for formal computational models of causal inference during event comprehension.

One prevailing theory characterizes causal inference as a process of probabilistic inference^13^. In this framework, people infer the causes of an event by performing Bayesian inference over hypothesis spaces generated by prior knowledge. Although this model has successfully explained human behavior in intuitive physics and early cognitive development^14–16^, it has not yet been generalized to the domain of event comprehension where events and causal links are continuously updated over time—which makes it a computationally intense problem. Another line of work has used variants of recurrent neural networks (RNNs) to directly simulate the event comprehension process. Studies have shown that the model segments continuous experience into meaningful events^17,18^, selectively encodes and retrieves memories at event boundaries^19^, and represents event schemas that give rise to structured event representations^20,21^. Yet, no event comprehension model thus far has examined the causal inference process.

Here, we introduce a model to test the idea that causal inference emerges from the continuous encoding and retrieval of episodic memories. By training the model to predict the next scene while “watching” a television episode, we examined whether the model would retrieve causally related past events to guide its predictions. Because exhaustive Bayesian inference is computationally costly, we considered an alternative possibility that organizing event representations based on their causal links may provide an efficient solution to the causal inference problem. We therefore hypothesized that, by selectively retrieving causally related memories, the model would learn to represent causally related events with similar patterns, organizing event representations based on their causal structure.

Our model is built on a gated recurrent unit (GRU) RNN to process dynamic inputs^22^. The RNN is augmented with an episodic memory (EM) buffer in which encoding and retrieval occur at every time step.

This design builds on recent memory-augmented neural networks that leverage an external memory buffer to store multiple memories and later retrieve them when long-term dependencies are needed^23–28^.

Encoding and retrieval are controlled by a key-value memory system that utilizes a self-attention mechanism in Transformers^29^. In this system, values represent memory contents whereas keys serve as memory addresses or indices^30^. This dissociation is analogous to a library search, where one uses an index (key) to locate a book (value). At each time step, a query surveys the memory buffer and assigns high attention weights to keys most similar to the query, but then retrieves values that are associated with those keys. In other words, the query searches through keys to find the ones with high pattern similarity, but the actual contents being retrieved are the associated values that hold orthogonal representations to the keys. By designing the model to use distinct representations for memory, we hypothesized that this would allow the system to retrieve memories based on higher-order event structure that goes beyond linear pattern similarity. The higher-order event structure was operationalized as the causal relationships scored by human raters in the previous study^12^.

This critically distinguishes the key-value framework from more commonly used content-addressable memory models, which retrieve memories by directly matching the current state to stored contents^25,26,31–33^. The key-value memory system has recently been proposed as a candidate mechanism of human memory and hippocampal function^30,34–37^ and has shown success in various cognitive tasks when integrated with an RNN^38,39^. To formally test its role in causal comprehension, we “lesioned” different components of the EM, such as keys, queries, or the entire EM buffer, to assess their respective contributions to event representations and memory retrieval.

The model’s internal representations and memory retrieval behavior were examined in relation to human behavioral and fMRI data collected by Song et al.^12^. In this empirical study, 36 participants watched a hard-to-understand television episode that was segmented into 48 events (∼52 s per event) and scrambled in time. Participants were asked to press an “aha” button whenever they understood something new about the narratives and were later asked to verbally explain their insights at those moments. Participants retrieved past events that were causally related with an ongoing event when explaining their insights. The study also found that the brain represents causally related events with similar neural patterns, suggesting that the brain formed event representations based on their causal structure. In our study, the model watched the same episode to match human experience. We asked if the model, like humans, retrieves past events that are causally related to the ongoing event. We further asked if the model represents events like the human brains do—by representing causally related events with similar patterns.

### A recurrent neural network with a key-value episodic memory buffer performs well in the next-scene prediction task and generalizes well to an unseen episode

We employed a GRU RNN augmented with a key-value EM buffer, hereby called the EM-GRU model (**Figure 1**). At each time step *t*, the model receives an input vector 𝑥_*t*_ which is transformed into three distinct representations: a hidden state ℎ_*t*_, a key 𝑘_*t*_, and a query 𝑞_*t*_. The hidden state ℎ_*t*_ is produced by a standard GRU update, based on the current input 𝑥_*t*_ and the previous hidden state ℎ_*t*−1_. This representation serves as a “value" in the episodic memory system and is stored in the EM buffer (EM[V]) at each time step. In parallel, the input 𝑥_*t*_ is linearly transformed into a key representation 𝑘_*t*_, which serves as a memory address and is also encoded in the EM (EM[K]). To retrieve relevant past information, a query 𝑞_*t*_(also linearly transformed from the input) surveys every stored key in EM[K]. High attention weights are given to selective past keys that are high in similarity with the query via the softmax function, following the self-attention mechanism used in Transformers^29^. Stored values (memory contents) associated with these keys (memory addresses) are retrieved as memory 𝑚_*t*_, computed as a weighted sum of past values with weights determined by the attention scores over the keys. Note that self-attention is calculated at the key space, but memory retrieval is happening at the value space. The ℎ_*t*_ representing the current input and 𝑚_*t*_ representing the retrieved memory are integrated via weighted summation: ℎ_*t*_ × 𝛼 + 𝑚_*t*_ × (1 − 𝛼), with 𝛼 set to 0.5. This integrated representation is then used to predict the next scene 𝑥_*t*+1_. The model was optimized for the next-scene prediction, with the goal of minimizing the loss between the predicted and observed 𝑥_*t*+1_. This objective function was motivated by a widely held view that people continuously generate predictions of upcoming events during comprehension^40–43^.

**Figure 1.**
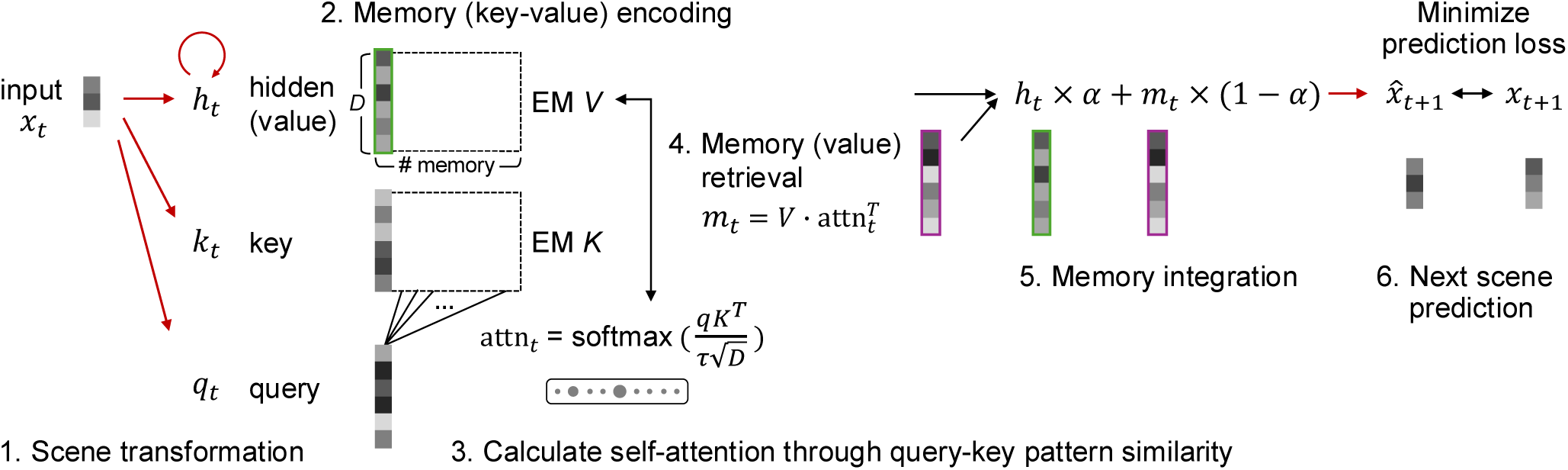
Model architecture. This figure illustrates a set of processes taking place at one time step. 1) *Scene transformation*. The model receives an input vector 𝑥_*t*_ at time *t* and transforms it into three distinct representations, a hidden state ℎ_*t*_, a key 𝑘_*t*_, and a query 𝑞_*t*_. Input-to-hidden transformation follows a nonlinear GRU RNN, and the hidden state corresponds to a value in the key-value system. Input-to-key and input-to-query mappings follow a linear transformation. 2) *Memory (key-value) encoding*. The hidden state and key vectors are encoded in the EM. The EM buffers are thus matrices in which the number of stored memories (rows) increases over time up to a predefined limit set by a hyperparameter. In our model, the number of memories corresponds to the number of scenes in episode 1, which means the model can store memories of all past scenes at test without forgetting. 3) *Calculate self-attention through query-key pattern similarity*. The query vector surveys all past keys stored in the EM and assigns attention weights based on the softmax function. The lower the 𝜏, the more selective the model becomes in assigning weights. In our model, 𝜏 was set to 0.1 to increase sparsity. Circles illustrate sparse attention weights given to past keys. See **Supplementary Figure S1** for results with varying 𝜏. 4) *Memory (value) retrieval*. Stored values in EM are summated, weighted by attention weights, which results in the retrieved memory 𝑚_*t*_. 5) *Memory integration*. The current scene representation ℎ_*t*_ is combined with the retrieved memory 𝑚_*t*_. 𝛼 is set to 0.5, such that the past and present representations are averaged. See **Supplementary Figure S1** for results with varying 𝛼. 6) *Next scene prediction*. The model learns a linear transformation from an integrated representation to the predicted next scene representation. The model parameters, indicated with red arrows, are optimized using gradient descent to minimize the loss between the predicted and observed next scenes. See ***Methods*** for details.

We asked whether the model, while watching a television episode, would learn the underlying event structure and retrieve relevant memories to guide comprehension. We used eighteen episodes from *This Is Us* Season 1, a character-focused television drama that follows the lives of members of a family, jumping back and forth in time from their early childhood to older adulthood. In turn, the narrative revolves around the same main characters, but at different stages of their lives. To represent the narratives, each episode was segmented into a sequence of ∼4-second scenes based on shot changes, which resulted in, on average, 614 ± 68 scenes comprising an episode. Visual and semantic contents of each scene were represented as a 512-dimensional embedding vector using a pretrained Contrastive Language Image Pretraining (CLIP) model^44^, which was reduced to the top 50 principal components. These embedding time series served as inputs to the model.

The model was trained to repeatedly watch episodes 2-18, during which model parameters were optimized through a gradient descent. We used these episodes because we wanted the model to acquire some general knowledge, such as how this television series was filmed and how characters in this series usually behave or interact with one another. We then fixed the learned model parameters and tested the model on the held-out episode 1. In Song et al.^12^, episode 1 was segmented into 48 events (mean 52 ± 12s) and presented in three different scrambled sequences to three groups of fMRI participants (N=12 per group).

For the model, the same stimulus was also presented in three different scrambled sequences at test, such that the results could be comparable to human participants’ neural and behavioral results. Note that temporal scrambling was at the coarse event-level (∼52 s per event), not the fine scene-level (∼4 s per scene), meaning both participants and the models watched scenes in their original order within each event, but only the event sequence was scrambled in time.

We compared the EM-GRU to four control models (**Figure 2**). First, to test the contribution of the self-attention mechanism, we shuffled the attention weights—after calculating them as in EM-GRU—so that the model retrieves random memories (*shuffled memory*; **Figure 2B**). This tested whether the EM-GRU’s attention to selective memories was meaningful and led to better performance than retrieving random ones. Second, to test the effect of having distinct representations for encoding and retrieval, we lesioned keys from the EM (*no key*; **Figure 2C**). In this variant, inputs were transformed into values and queries, and the query directly searched stored values based on pattern similarity, using the same attention calculation as EM-GRU. This provided a comparison between the key-value mechanism and content-addressable memory retrieval^45^. Third, to test the effect of removing both keys and queries, we implemented a model frequently used in prior literature (*no key-query*; **Figure 2D**)^25,26,31–33^. In this design, the current hidden state acts like the query, searching past hidden states to find those with high pattern similarities. Notably, there was no additional transformation beyond the input-to-hidden state mapping, meaning pattern similarity was computed within the same representational space. Finally, to assess the benefit of having an episodic memory buffer with the RNN, we trained a standard GRU RNN without the external EM (*no EM*; F**igure 2E**). This baseline model did not have keys, queries, or memory encoding and retrieval processes that took place at every time step.

**Figure 2.**
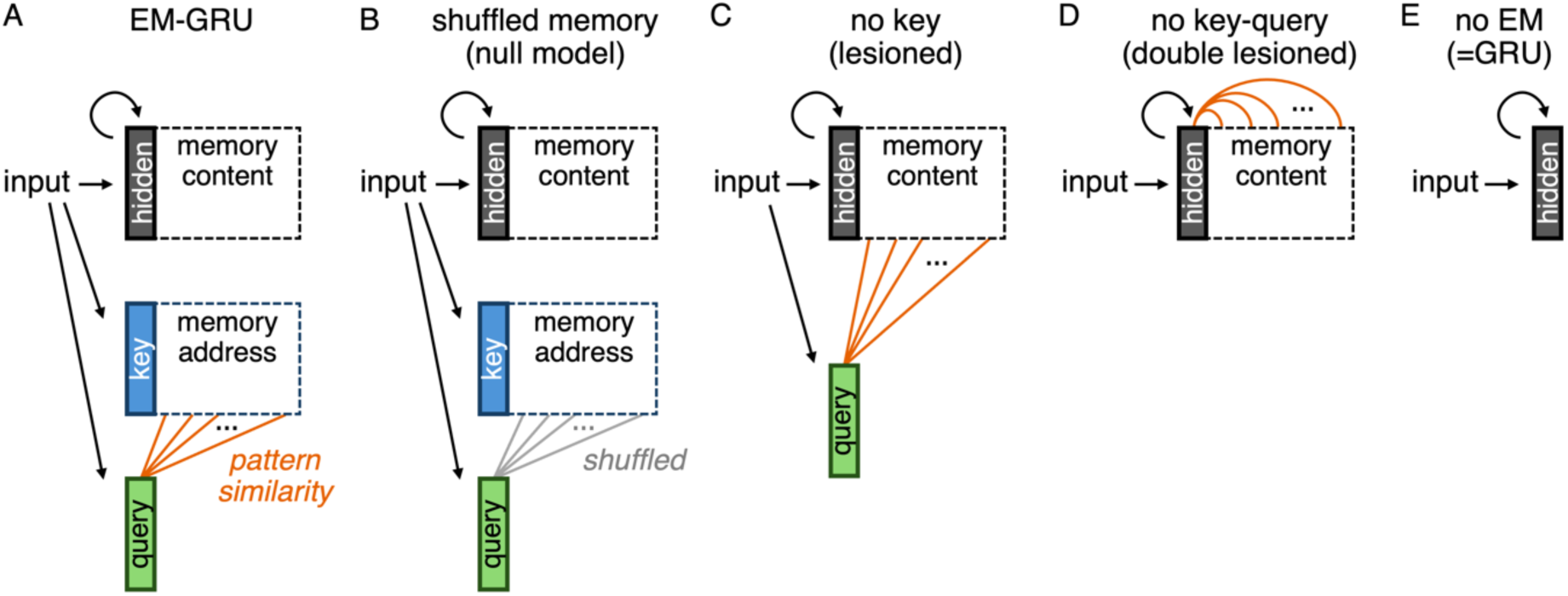
EM-GRU and comparison models of EM variants. **(A)** EM-GRU model. A simplified visualization of our main model in Figure 1, focused on input transformation and memory retrieval based on pattern similarity. **(B)** *Shuffled memory* model. The architecture is the same as EM-GRU, but we shuffled attention weights calculated between a query and keys, so that the model retrieves random memories. This serves as a null model that tests whether memories retrieved by the model are informative for task performance and event representations. **(C)** *No key* model. The model does not have keys that address memory contents. Thus, pattern similarity is calculated directly between a query and hidden states. This model tests the usefulness of dissociating representations used for memory contents and addresses. **(D)** *No key-query* model. The model has neither a key nor a query. Instead, the hidden state of the current moment acts like a query and searches through past hidden states stored in the EM. This design follows the conventional approach in memory research, retrieving information by matching the current state directly to past stored states. **(E)** *No EM (GRU)* model. The model is a GRU RNN that is not augmented with EM. It is designed to test the utility of retrieving memories from the distant past in addition to the recurrence implemented by the GRU. Black arrows indicate learnable model parameters. Orange lines indicate attention calculation based on pattern similarity, such that memories that are assigned high attention weights due to high pattern similarity are selectively retrieved.

**Figure 3** shows EM-GRU and the control models’ next-scene prediction performances over the course of multiple training iterations. All five models learned well (solid lines in **Figure 3A**) and generalized well (**Figure 3B**; **Supplementary Figure S2**). All models learned rapidly up to near 10th training iteration, after which the loss decreased more gradually. Generalization loss at test was minimal near the 35th training iteration and rose after that. Thus, we stopped the training at the 50th iteration and report selective findings from the 35th iteration in this paper. The *no EM (GRU)* model learned the fastest among the five but eventually converged to a similar level of performance as the EM-GRU. The *shuffled memory* model performed the worst among the five models.

**Figure 3.**
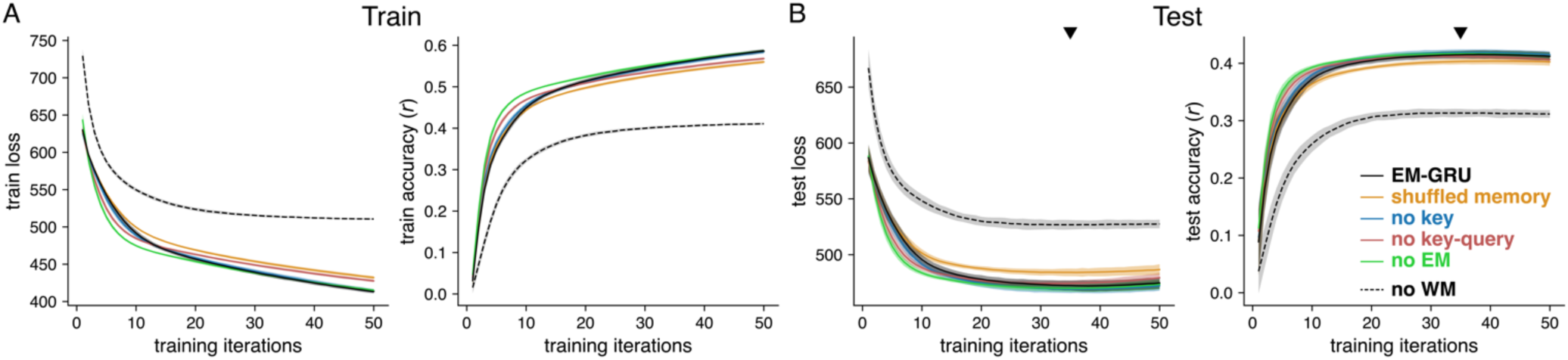
Model performance at train and test. **(A)** Train performance. One training iteration includes 17 rounds of parameter updates as the model watches episodes 2 to 18. **(B)** Test performance. The model was tested on the held-out episode 1 after different rounds of training iterations. Loss indicates the mean squared error between the predicted and observed next scene vectors, averaged across time. Accuracy indicates Pearson’s correlation between the predicted and observed next scene vectors, averaged across time. Black lines indicate EM-GRU results, and the colored lines indicate results from the comparison models of EM variants. Black dashed lines indicate results of the no working memory (WM) model in which the GRU recurrence was lesioned. Lines indicate the mean and error bars indicate 95% confidence intervals across 20 repetitions using different random seeds. The triangle points to the 35th training iteration where the loss is minimized at test.

All of these models, which were EM variants, performed comparably on the next scene prediction task. We asked whether they performed comparably because the prediction task was mainly supported by the GRU rather than the EM. To test this, we lesioned the recurrence in the GRU which was designed to implement working memory (WM; *no WM* model). Removing the recurrence significantly reduced performance (dashed lines in **Figure 3**; **Supplementary Figure S3**). This suggests that the EM plays a relatively minor role in the next-scene prediction task. While next-scene prediction was an explicit objective function used to train the models, the primary focus of this study is on the emergent property afforded by the EM— specifically, how the model organizes events and retrieves episodic memories during comprehension. Therefore, the following sections focus on properties that naturally emerged during training, beyond what have explicitly been trained.

### The model retrieves memories similar to those retrieved by humans

The EM-GRU and the control models all performed and generalized well on the next-scene prediction task, which was explicitly optimized in training. More importantly, we asked whether the model retrieved memories that are meaningful for comprehension. To ask this, we compared memories retrieved by the model to memories retrieved by human participants as they both watched the same sitcom episode.

Human memory retrieval was directly coded from participants’ verbal responses made during the experiment. In Song et al.^12^, more than 40% of people’s insight explanations included mentions of past events. If an event A was retrieved at an insight during event B, +1 was coded in the events A-B pair in the memory retrieval matrix. The memory retrieval matrices created for all participants were averaged (**Figure 4A** *Left*).

**Figure 4.**
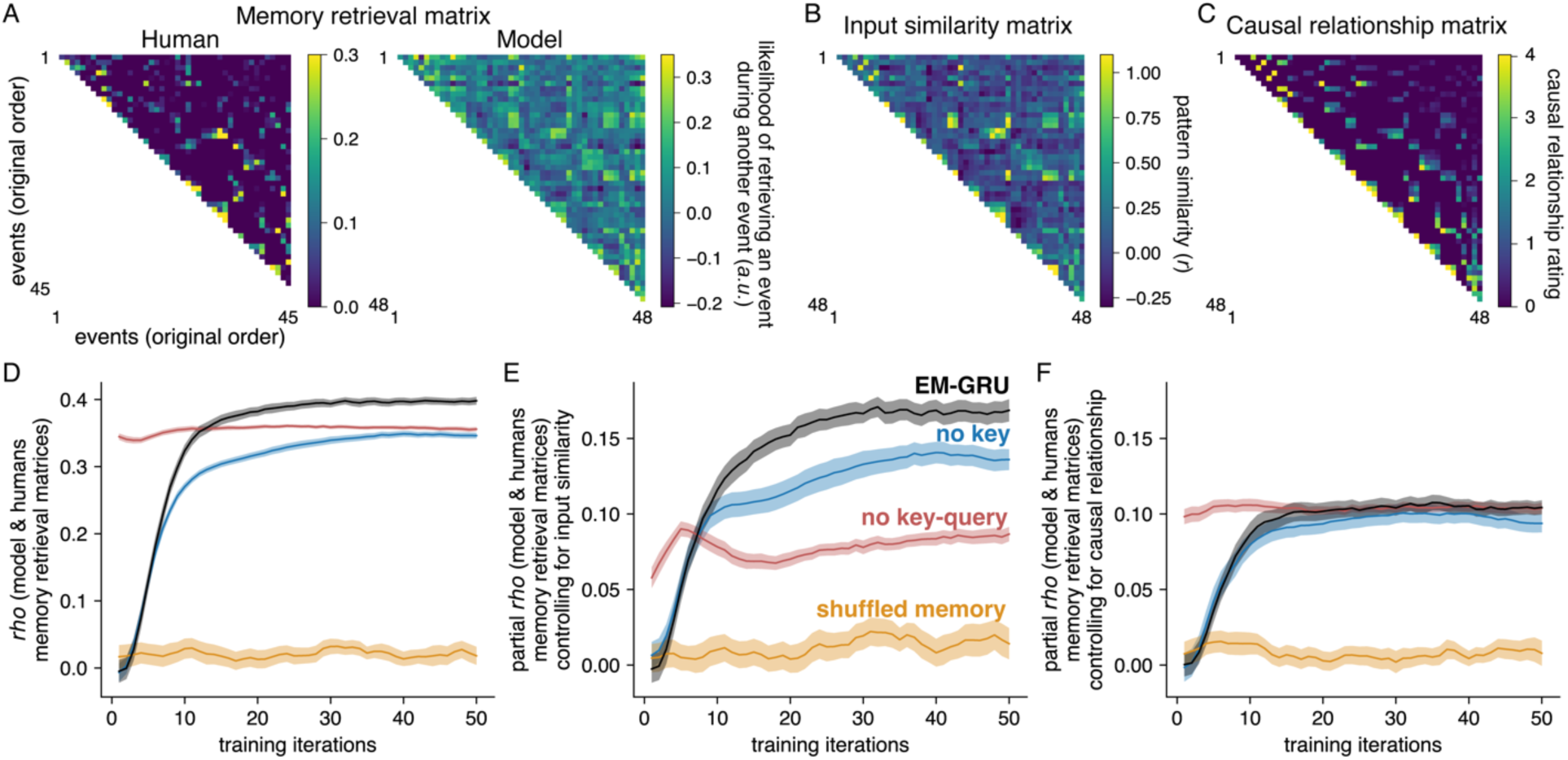
Human and model memory retrieval. **(A)** Memory retrieval matrices of the model and human participants. (*Left*) Humans’ memory retrieval matrix from Song et al.^12^ based on participants’ verbal explanations of their insight moments. If an event was retrieved when explaining an insight during another event, +1 was noted for that event pair. Memory retrieval matrices of 32 participants assigned to three different scrambled order groups were averaged. Events 46 to 48 were not included in analyses as in Song et al.^12^. (*Right*) Because the likelihood of retrieving one event during another event was determined by pattern similarity between the query and the key, the model’s memory retrieval was represented by the query-key pattern similarity. Memory retrieval matrices created for three different scrambled sequences and 20 different random seeds were averaged. The model’s memory retrieval matrix of 35th training iteration is visualized. **(B)** Input similarity matrix. The CLIP embeddings of all scenes corresponding to one event was averaged to represent an input pattern for that event. We calculated an event-by-event input pattern similarity, representing the degrees of semantic and perceptual feature similarities of all event pairs, extracted from CLIP. **(C)** Causal relationship matrix. Three raters rated how causally related every event pair is, on a scale of 0 to 4. The three ratings were averaged to create a causal relationship matrix in Song et al.^12^. **(D)** Relating the model’s memory retrieval with humans’ memory retrieval. The model and humans’ memory retrieval matrices were correlated across different training iterations. High correlation between the two matrices indicates that the model retrieved similar event memories as humans did. **(E)** Relating the model’s memory retrieval with humans’ memory retrieval, while controlling for input pattern similarity in **B**. **(F)** Relating the model’s memory retrieval with humans’ memory retrieval, while controlling for causal relationships in **C**. Note that the *no EM (GRU)* model was not analyzed, as this model does not retrieve memories. Lines indicate the mean and error bars indicate the standard error of the mean across 20 repetitions using different random seeds.

Unlike the human memory retrieval matrix, which reflected selective retrieval at insight moments, the model’s memory retrieval matrix reflected retrieval occurring at every time step throughout comprehension. Our self-attention mechanism was based on pattern similarity between the current query and past keys, such that keys with higher pattern similarity with the query received higher attention weights. To assess whether one event was more or less likely to be retrieved by another event, we averaged the query and key representation patterns across all scenes corresponding to each of the 48 events. We then computed query-key pattern similarities for all event pairs, generating an event-by-event memory retrieval matrix of the model (**Figure 4A** *Right*). In control models that did not have queries or keys, the representations serving those roles were instead used to compute memory retrieval matrices (i.e., query-value similarity for the *no key* model and value-value similarity for the *no key-query* model).

Over the course of training, the correlation between the model’s and humans’ memory retrieval matrices increased up to near 𝜌 = 0.4 for EM-GRU (𝜌 = 0.397 ± 0.032 at training iteration = 35; **Figure 4D**). This indicates that the model became increasingly human-like in terms of which memories it selected for retrieval. The noise ceiling was 𝜌 = 0.647, estimated as the average split-half correlation across the 32 participants’ memory retrieval matrices. The resemblance between the EM-GRU model’s and humans’ retrieval matrices was stronger than that of the *no key* model (𝜌 = 0.345 ± 0.026) and the *shuffled memory* model (𝜌 = 0.022 ± 0.042). These results suggest that including keys, and thus having separate representations for memory contents and addresses, was useful for memory retrieval. They also suggest that the self-attention mechanism selected memories that meaningfully mapped onto human behavior.

Notably, the *no key-query* model, which directly retrieves past hidden states based on their similarity with the current hidden state, immediately produced human-like retrieval even before training. We reasoned that this could be due to an intrinsic correlation between memory retrieval and the model inputs’ pattern similarity (i.e., CLIP embedding similarity), which reflects semantic and perceptual similarities between events (𝜌 = 0.472, *p* = 5e-56; **Figure 4B**). To account for this, we controlled for the input pattern similarity matrix when correlating the model’s and humans’ memory retrieval matrices. Even after this control, the EM-GRU model showed the highest correlation with humans compared to the control models (partial 𝜌 = 0.169 ± 0.032, with the noise ceiling for the partial 𝜌 = 0.558; **Figure 4E**). As expected, the correlation of the *no key-query* model dropped substantially (partial 𝜌 = 0.081 ± 0.019), to a level below that of the *no key* model (partial 𝜌 = 0.137 ± 0.036). The EM-GRU model significantly outperformed all control models (paired *t*-tests: *t*(19) > 3.85, FDR-corrected *p* < .002, Cohen’s *d* > .92). Together, these findings suggest that the EM-GRU model retrieved memories most closely aligned with those of humans watching the same movie.

### Causal event structure accounts for the similarity between model and human memory retrieval

How did the EM-GRU learn to retrieve memories like humans? Song et al.^12^ found that participants retrieved events that were causally related to the current event, which was based on a theory that people retrieve causally related past events during comprehension of an ongoing event^2–5,7,9–11,46^. To test this, the study rated the degree of causal relationship between every event pair to create a causal relationship matrix (**Figure 4C**; **Supplementary Text S1**). They found that this causal relationship matrix explained significant variance in the memory retrieval matrix, beyond what could be explained by semantic and perceptual similarities. Based on this finding, we hypothesized that the model would have learned to retrieve like human participants by having captured causal relationships between narrative events.

To test this, when correlating the model’s memory retrieval matrix with that of humans’, we controlled for causal relationships, similar to how we controlled for input similarity in **Figure 4E**. The partial correlation value of the EM-GRU was lower than that of **Figure 4E** (𝜌 = 0.108 ± 0.023 at training iteration = 35), and importantly, they were comparable to *no key* (𝜌 = 0.100 ± 0.028) and *no key-query* (𝜌 = 0.104 ± 0.013) models (**Figure 4F**). This suggests that the EM-GRU may have retrieved memories like human participants because it captured causal relationships between events.

But how could an artificial agent capture higher-order relational structure beyond what was explicitly provided as input? In addition to assessing the role of causal relationships in explaining the similarity between model and human retrieval, we assessed their role in directly explaining the model’s memory retrieval.

We first designed a regression model predicting the humans’ memory retrieval matrix (**Figure 4A** *left*) from input similarity (**Figure 4B**) and causal relationship (**Figure 4C**). Causal relationship explained significantly more variance (*t*(987) = 21.802, *p* < .0001) than input similarity (*t* = 4.661, *p* < .0001), and including causal relationship as an independent variable significantly improved model fit relative to a model without it (F(1, 987) = 475.307, *p* = 2.5e-86).

In contrast, this pattern flipped when predicting the model’s memory retrieval matrix (**Figure 4A** *right*). Input similarity was a stronger predictor of the model’s memory retrieval (*t*(987) = 28.438, *p* < .0001) than causal relationship (*t* = 1.755, *p* = .080). Including causal relationship as an independent variable did not significantly improve model fit relative to a model without it (F(1, 987) = 3.079, *p* = .080). These results suggest that, whereas human memory retrieval is primarily explained by causal relationships, the model’s memory retrieval is more explained by semantic and perceptual feature similarities that are explicitly provided as inputs.

Together, these findings highlight nuanced implications for the EM-GRU. Whereas human memory retrieval primarily relies on causal structure, the EM-GRU relies more heavily on semantic and perceptual features. This suggests that the EM-GRU is an imperfect model of human memory. Nevertheless, the EM-GRU’s memory retrieval is more similar to human retrieval than those of the control models only before controlling for causal relationships (**Figure 4E and 4F**). Once the variance explained by causal structure is removed, this advantage disappears, indicating that the human-like behavior of the EM-GRU may be driven by its alignment with causal structure that goes beyond input similarity.

### The model represents causally related events with similar patterns

The EM-GRU learned to retrieve memories like humans during movie-watching. We then asked whether, through retrieving these memories, the model also came to represent causally related events with more similar patterns. Song et al.^12^ showed that causally related events are represented by similar brain patterns, beyond semantic and perceptual similarities. Here, we replicated this analysis, but by replacing brain representations with the model’s internal representations.

At test, we extracted the time series of the GRU hidden states ℎ_𝑇_. Representation patterns of all scenes corresponding to each of the 48 events were averaged to represent that one event. By taking correlations of all pairs of these 48 events, we created an event-by-event representation similarity matrix (RSM; **Figure 5A**). The RSM was then correlated with the episode’s causal relationship matrix in **Figure 4C**.

**Figure 5.**
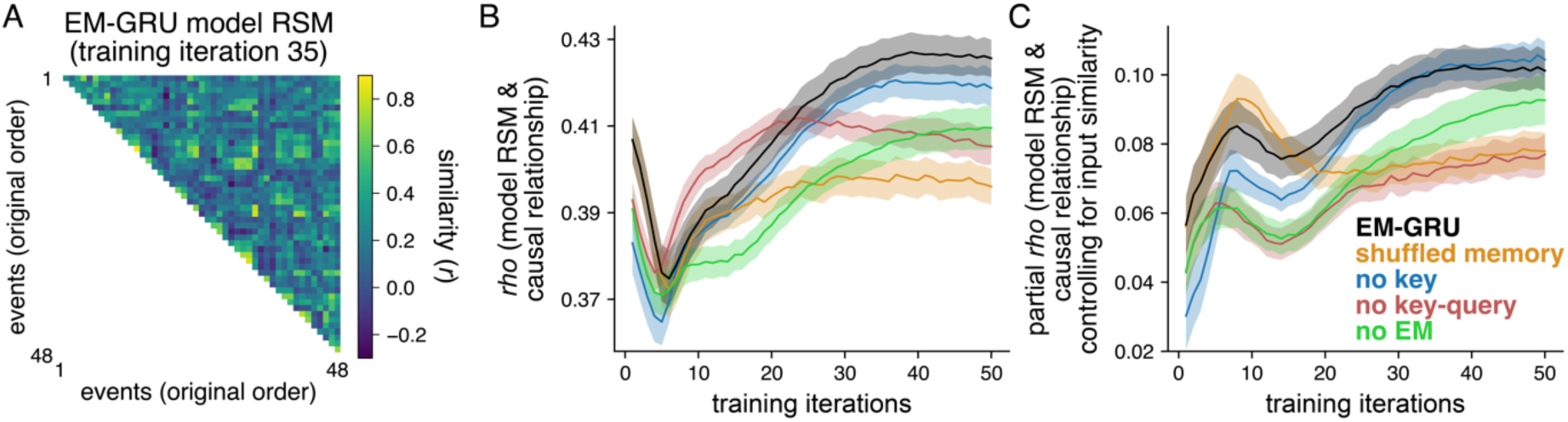
Models’ representation of causal event structure. **(A)** Event-by-event model representation similarity. Upon training the model 35 times and testing it on episode 1, we extracted the representational time series of the hidden state, or value, ℎ_𝑇_. Scene patterns pertaining to an event were averaged, and the resulting patterns representing 48 events were correlated in pairs. This resulted in an event-by-event representation similarity matrix (RSM). RSMs of 20 different random seeds were averaged for visualization. **(B)** Relating model representation similarity with causal relationships. The model RSMs were correlated with the human-rated causal relationship matrix in Figure 4C across different training iterations. High correlation between the two matrices indicates that the model represents causally related events with similar patterns. **(C)** Relating model representation similarity with causal relationships, while controlling for input pattern similarity in Figure 4B. Lines indicate the mean and error bars indicate the standard error of the mean across 20 repetitions using different random seeds.

Across training, the EM-GRU increasingly reflected events’ causal relationship structure in its representations (**Figure 5B**). The EM-GRU model RSM consistently showed stronger correlation with the causal relationship matrix than comparison models. In other words, the model learned to represent causally related events with similar patterns, whereas this trend was weaker in control models. Although EM-GRU (𝜌 = 0.425 ± 0.026) showed numerically higher correlations than the *no key* model (𝜌 = 0.419 ± 0.020), the two were statistically not significant in how much their RSMs captured causal relationship structure (paired *t*-test: *t*(19) = 1.028, FDR-corrected *p* = 0.317, Cohen’s *d* = .331; whereas EM-GRU compared to other models: *t*(19) > 2.87, FDR-corrected *p* < .02, Cohen’s *d* > .80). This suggests that a model where queries directly search through memory contents without the keys has sufficiently captured the causal relationship structure as the EM-GRU.

This pattern of results persisted after controlling for input pattern similarity (**Figure 5C**). Although partial correlation values were numerically smaller (partial 𝜌 = 0.101 ± 0.029), they remained significantly above chance, as confirmed by comparison with a null distribution generated by shuffling model RSMs 1,000 times (*z* = 15.295, *p* < 0.001). (Note that the partial 𝜌 does not start at 0 at the start of training, because the GRU nonlinearly transforms the inputs, which may have introduced variance that cannot be fully controlled for. Furthermore, a sudden increase in partial *rho* values near the 8th training iteration across all models appears to be driven by concurrent decreases in correlations between the model RSM and both the causal relationship (**Figure 5B**) and input similarity (**Supplementary Figure S4**), rather than reflecting a meaningful change in representational structure.) Together, the findings indicate that, while the difference to control models are nuanced and the partial correlation value is numerically small, the EM-GRU represents causally related events with more similar representation patterns, mirroring how the human brain represents causally related events.

### The model’s event representation resembles that of the human brain

Finally, we asked if the model represents narrative events similar to how the human brain represents the same events during fMRI. For each participant in Song et al.^12^, we extracted voxel-wise BOLD activity time series for each of the 200 cortical^47^ and 32 subcortical regions^48^ and averaged the time series per event to extract voxel activity patterns representing 48 events. For every brain region, we calculated an event-by-event RSM, estimating how similar the voxel activity patterns are for every pair of 48 events. We correlated the brain RSM to the model RSM (**Figure 5A**), to ask if the model’s event representation resembles that of the functional brain.

Among the 232 areas tested, we found that most brain areas’ RSMs positively and significantly correlated with the EM-GRU model RSMs, even upon controlling for input pattern similarity (at training iterations = 35), compared to chance distributions where brain RSMs were randomly shuffled (1,000 iterations).

Specifically, the representation of the model’s hidden states resembled 207 out of 232 brain regions’ representational patterns significantly above chance (FDR-corrected *p* < .05) (**Figure 6A**). These suggest that the model’s representations resemble the representational patterns observed in the majority of brain areas.

**Figure 6.**
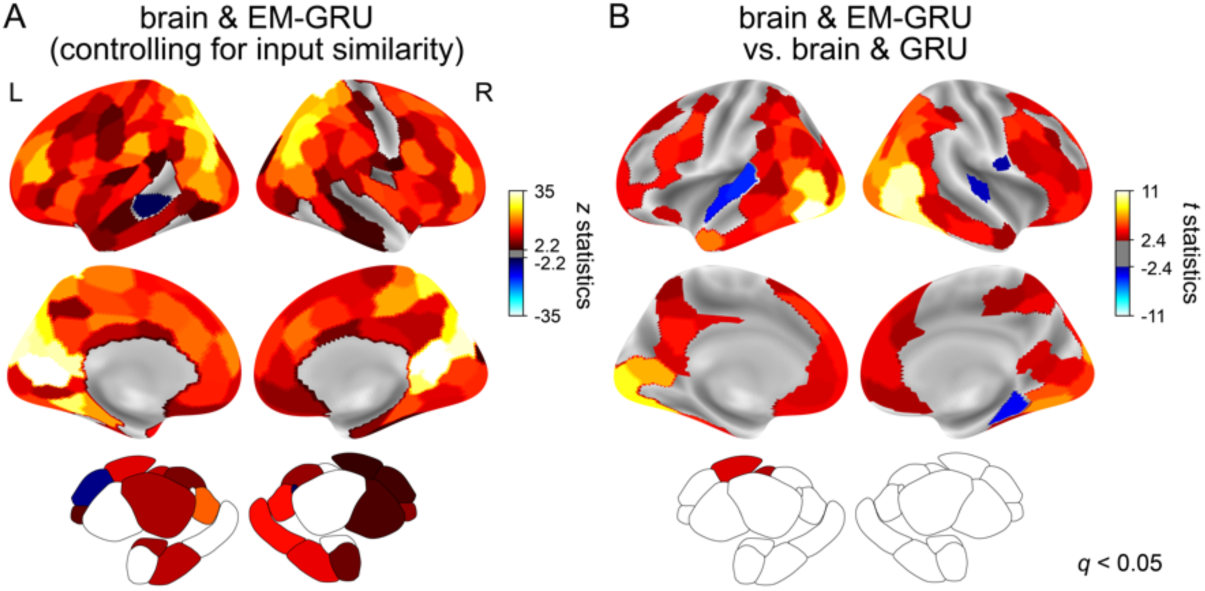
Model and human brain’s representation similarity. **(A)** Comparison between the EM-GRU model representation similarity matrix (RSM) with the human brains’ RSMs. The EM-GRU model RSM was correlated with the voxel pattern RSMs of each brain area of 33 participants (controlling for input pattern similarity), and compared to a chance distribution in which brain RSMs were randomly shuffled. Colors denote z statistics. **(B)** Paired comparison between the EM-GRU and GRU model RSMs in how much they resemble brain RSMs. The EM-GRU and GRU models’ RSMs were correlated with the voxel pattern RSMs of each brain area of 33 participants. These 33 correlation values of the EM-GRU and GRU models were compared using paired *t*-tests, repeatedly across 232 brain regions. Colors denote *t* statistics. Red-yellow colors indicate that the region’s representation was more similar to the EM-GRU, whereas blue-cyan colors indicate that the region’s representation was more similar to the GRU. Colored are brain regions that hold statistical significance (FDR-corrected *p* < 0.05, corrected for 232 brain regions). The figure shows the lateral (*Top*) and medial (*Middle*) visualizations of the cortex and the lateral visualization of the subcortex (*Bottom*). See **Supplementary Figure S6** for an unthresholded result.

Importantly, we asked whether the inclusion of the EM buffer boosted the model’s representational similarity with the brain. We tested whether the EM-GRU model RSM is more similar to brain RSMs, compared to the GRU model RSM. We selected the GRU as the comparison model because this model does not have an EM buffer (but see **Supplementary Figure S5** for comparisons with different control models). Many brain regions—except for the sensorimotor areas—exhibited significantly higher similarity with the EM-GRU’s hidden states compared to the GRU’s hidden states (paired *t*-tests, FDR-corrected *p* < .05) (**Figure 6B**). This suggests that the EM improved the RNN’s hidden state representations, making them not only better reflect causal event structure but also more closely match the human brain’s representational patterns.

### The model transforms inputs into separate value, key, and query representations

We designed the EM-GRU to use distinct representations for values, keys, and queries. To examine how these representations evolved over training, we measured similarities between values, keys, and queries across training iterations (**Supplementary Figure S7**). Correlations between value–query and value–key representations remained near zero, suggesting that the model maintained orthogonal representations for memory contents (values) and memory addresses (keys and queries). The query surveys the keys to select memories based on pattern similarity, but retrieval occurs in the value space, which stores memory contents that are orthogonal to the keys. This separation allows retrieved memories to differ from the addressing representation, enabling the model to represent relationships between events beyond simple similarity within a single representation space.

In contrast, key–query similarity increased over training and stabilized around 𝜌 = 0.3 (𝜌 = 0.277 ± 0.015 at training iteration = 35; **Supplementary Figure S7**). Even so, this model with a moderate degree of key– query similarity retrieved memories more like humans compared to a model in which keys and queries were forced to be identical (paired *t*-test: *t*(19) = 8.227, *p* = 1e-07; partial 𝜌 = 0.092 ± 0.034, compared to the original EM-GRU of partial 𝜌 = 0.169 ± 0.032). When keys and queries are identical, the attention mechanism (𝑞𝐾^𝑇^) simply reduces to a dot-product similarity. In contrast, separating the key and query projections allows the attention mechanism to depend on two independently learned transformations of the input (i.e., a bilinear function), enabling the model to capture more flexible and potentially asymmetric relationships between current scenes and stored memories. Therefore, these findings suggest that maintaining three separate representations for narrative events may be a key ingredient enabling the EM-GRU to retrieve memories and represent events in a human-like manner.

## Discussion

In this study, we introduced an RNN augmented with a key-value EM buffer that continuously encodes and retrieves memories to comprehend an ongoing event. We found that the model retrieves memories that are comparable to what human participants retrieve and represents causally related events with similar patterns. Moreover, its representational structure aligns more closely with human brain patterns than that of the control models. These findings highlight EM-GRU as a candidate model of human memory during event comprehension and suggest a computational mechanism by which we remember and reason about ongoing naturalistic events.

We developed EM-GRU to understand how causally related past events are selectively retrieved during event comprehension. Inspired by and building on recent memory models^19,30,38,39^, the EM-GRU is comprised of two parts: the GRU, whose recurrence represents working memory^38,39^, and the key-value EM buffer, which grants full access to memories of the distant past. The model was not explicitly trained to perform causal inference but was optimized for next-scene prediction. The GRU was found to be largely responsible for the next-scene prediction task, given that models of EM variants showed comparable task performance while lesioning the recurrence significantly decreased it. Interestingly, when we enforced the model to use retrieved memories when generating predictions, the EM-GRU retrieved memories closely aligned with those of human participants. Moreover, the hidden states of the GRU, when augmented with the key-value EM, represented causally related events with similar patterns more so than the control models. These findings support the view that causal inference during comprehension emerges from continuous memory encoding and retrieval. The GRU mainly performs next-scene prediction, while the EM specializes in retrieving relevant memories and helps the GRU to represent events based on causal relationships.

What features of the model allowed it to capture causal dependencies and assign high attention weights to causally related past events? Two features of the EM-GRU appear critical for the observed correspondence with the human brain and behavior. First, the model uses two distinct representations for memory: values representing memory contents and keys representing memory addresses. A query surveys the keys based on pattern similarity, but retrieval occurs in the value space whose representations are orthogonal to the keys. That is, memory retrieval involves a two-step process: addressing in key space, then retrieving values associated with the keys. Furthermore, even during memory addressing in key space, the model computes attention using separate key and query representations, so that pattern similarity is not constrained to a simple dot product between movie scene representations. We hypothesized that using these diverse representational spaces allows the model to retrieve memories based on relationships that extend beyond linear pattern similarity, thereby capturing underlying causal dependencies between events. These features distinguish the EM-GRU from classic frameworks of episodic memory—such as the Hopfield network^31^, the temporal context model^32,33^, and a range of EM-augmented RNN models^25,26^—which use a single representational scheme to directly store internal representations in episodic memory and later retrieve those with high pattern similarity to the current state.

Second, EM-GRU includes two notable integration processes. One involves integrating past memories based on attention weights calculated from the self-attention mechanism. This is motivated by a view that people retrieve not just one memory, but a blend of multiple memories that are assigned different weights^49^. Yet, there is a sweet spot—the model performs better if it is attending to a few selective memories by assigning sparse attention weights compared to broadly summing up arrays of memories using diffuse attention weights (**Supplementary Figure S1**)^50–52^. The second integration combines the current representation (ℎ_*t*_) with the retrieved memory (𝑚_*t*_) to predict the upcoming scene. This integration of the past and the present did not improve accuracy in the next-step prediction task (**Figure 3**). However, it was critical for emergent causal reasoning and contributed significantly to the model’s alignment with human behavior (**Supplementary Figure S1**).

Overall, the model performance and correlation values reported in this manuscript are not high. The aim of this paper was not to establish a performance benchmark or to achieve state-of-the-art accuracy. Our aim was to use models to test cognitive and neuroscientific hypotheses about how the brain organizes and retrieves causally related memories during comprehension. Therefore, we designed a model with clear hypotheses and compared it against control models in which the component of interest (i.e., EM) was lesioned or systematically varied, rather than benchmarking it against pre-existing machine learning models. We therefore orient readers’ attention to the relative differences between models, rather than to their absolute performance values.

The EM-GRU is a normative model of cognition that does not simulate neurobiological circuits. Although we have seen alignment between the EM-GRU and the human brain’s representation patterns, this simply implies that the EM-GRU represents events in meaningful ways. Future work can continue to develop this model in relation to neurobiological circuits relevant to memory.

Furthermore, the current study is limited in that it is based on one episode of a television series. This choice was due to the availability of rich, pre-collected data that used this limited set of stimuli. The generalizability of the model should be examined in future work, potentially using a broader range of naturalistic stimuli.

In sum, the EM-GRU model, though not explicitly trained for causal inference, spontaneously developed internal representations that reflect causal relationships between events and exhibited memory retrieval behavior that resemble those of humans. These findings provide insights into the computational underpinnings of how people reason about causal relationships between events through memory encoding and retrieval.

## Data availability

Raw and processed fMRI data are shared in OpenNeuro, https://openneuro.org/datasets/ds005658^12^. Behavioral data are shared in GitHub, https://github.com/hyssong/memoryaha^12^.

## Code availability

Model and analysis codes and processed behavioral data are openly available in: https://github.com/hyssong/memorymodel.

## Acknowledgment

We thank Randall O’Reilly for insightful guidance during the early stages of model design. We thank JeongJun Park with conceptualization and helpful discussions and comments on the manuscript. We thank the two anonymous reviewers for their tremendous help in improving the manuscript. The research was supported by the McDonnell Center for Systems Neuroscience and the McDonnell Center for Cellular and Molecular Neurobiology at Washington University in St. Louis (HS).

## Competing interests

The authors declare no competing interests.

## Author contributions

HS: Conceptualization, Data curation, Formal analysis, Funding acquisition, Investigation, Methodology, Project administration, Visualization, Writing – original draft, Writing – review & editing

QL: Conceptualization, Investigation, Methodology, Supervision, Writing – review & editing TTN: Conceptualization, Investigation, Writing – review & editing

JC: Conceptualization, Investigation

YCL: Conceptualization, Investigation, Writing – review & editing MDR: Conceptualization, Investigation, Writing – review & editing SC: Conceptualization, Investigation

JMZ: Conceptualization, Investigation, Supervision, Writing – review & editing

## Methods

### Stimuli

Eighteen episodes of a television series, *This is Us* Season 1 (2016, directed by Requa, J. & Ficarra, G. and written by Fogelman, D.) were used as stimuli. Episodes 2-18 were used as training data (mean 41m 37s, ranging from 40m 16s to 43m 31s) and episode 1 (41m 40s) was used as test data. During test, the episode was segmented into 48 events (mean 52 ± 12s, ranging from 8s to 1m 37s) and ordered in a temporally scrambled sequence. Temporal scrambling followed the manipulation of Song et al.^12^, with the goal of relating model results to human behavioral and fMRI data. In Song et al.^12^, 36 fMRI participants were assigned to one of three groups where they watched the same events but in a different sequence. Each event was positioned at either the beginning, middle, or end of the sequence in the three groups, so that the order effect could be counterbalanced when averaging across groups. Episodes 2-18 were used in training without any manipulation to the video. Yet, the director has originally filmed the narrative to jump back and forth in time, switching between past, present, and future narratives of multiple characters.

To finely segment the episodes into multiple scenes with minimal autocorrelation between consecutive scenes, we applied an automated scene segmentation algorithm (PySceneDetect) that cuts the episode into multiple scenes based on automatically-detected camera shot changes. This resulted in on average 614 ± 68 scenes per episode (ranging from 523 to 751), with each scene lasting for 4.07 ± 4.78s (ranging from 0.13s to 182.68s).

To extract visual and semantic contents of each scene and transform it into a scene embedding, we used a pre-trained CLIP model^44^, which has often been used in previous fMRI studies^53,54^. We applied the CLIP model to the frames of each scene, extracting 512-dimensional embeddings representing each visual frame. Multiple frames’ embedding vectors were averaged to represent one scene, and the resulting embedding time series of hundreds of scenes represented the unfolding narrative of an episode. Upon concatenating the train data, we *z*-normalized each embedding feature across time, of which mean and standard deviation were applied to normalize the test data.

To constrain the number of model parameters, we applied a principal component analysis to the training data to reduce the number of dimensions to 50, which were again *z*-normalized across time per feature. We transformed the test data to the trained principal component space, where we again applied the mean and standard deviation to normalize the test data. These dimensionality-reduced embedding time series were used as inputs to the model, where the input at time *t* (𝑥_*t*_) corresponded to the embedding vector of a scene. Consecutive scene embeddings were significantly correlated (episode 1: *r* = 0.279 ± 0.272) compared to a chance distribution where embeddings were randomly shuffled across time (*z* = 28.261, *p* < .001). We considered this as an intrinsic characteristic of naturalistic movies because nearby scenes occur at similar place and time, with the appearance of overlapping characters.

To create an event-by-event input pattern similarity matrix, we averaged CLIP embedding vectors of all scenes that correspond to each of the 48 events to have one input pattern representing each event. Pearson’s correlation was calculated for all pairs of input patterns.

Audio information was not included as input. One reason was that 22.76% of the scenes in episode 1 contained no utterances. When transcribing characters’ conversations using WhisperX^55^, we found that the average number of words per scene was 5.655 ± 6.961, with a median of 3. More than half of the scenes (57.72%) included three or fewer spoken words, indicating that dialogue was sparse by the nature of this television episode. Another reason for excluding audio information was to make the model generalizable. If the model performs well using only visual information, it could potentially be applied to a wider range of naturalistic videos, regardless of whether audio is present. Therefore, we chose to report findings with the limited but generalizable embedding approach.

### Model

The core architecture of the model is a GRU RNN ^22^. At each time step *t*, the CLIP embedding 𝑥_*t*_ ∈ ℝ^𝑑^ (𝑑 = 50) was transformed into a 6𝑑-dimensional vector, decomposed into 𝑖_𝑟_, 𝑖_𝑧_, 𝑖_𝑛_ ∈ ℝ^2𝑑^, via the parameters 𝑾_𝒙_ ∈ ℝ^6𝑑×𝑑^ and 𝒃_𝒙_ ∈ ℝ^6𝑑^. The previous hidden state ℎ_*t*−1_ ∈ ℝ^2𝑑^ was likewise transformed into a 6𝑑-dimensional vector, decomposed into ℎ_𝑟_, ℎ_𝑧_, ℎ_𝑛_ ∈ ℝ^2𝑑^, using parameters 𝑾_𝒉_ ∈ ℝ^6𝑑×2𝑑^ and 𝒃_𝒉_ ∈ ℝ^6𝑑^. The reset gate 𝑟_*t*_ and update gate 𝑧_*t*_ were then computed to control how much of the previous hidden state was carried forward versus updated. The new hidden state ℎ_*t*_ was obtained as a transformed, weighted combination of 𝑥_*t*_ and ℎ_*t*−1_.

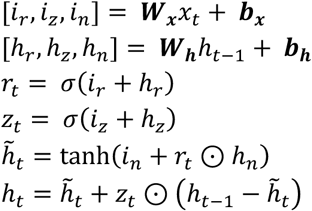

We augmented this RNN with an EM buffer, where an RNN can write to (i.e., encoding) and read from (i.e., retrieval) an external memory storage. Inputs are linearly transformed into keys (𝑘_*t*_ ∈ ℝ^2𝑑^) that represent memory addresses and are encoded in the EM, and queries (𝑞_*t*_ ∈ ℝ^2𝑑^) that match the keys to calculate self-attention based on similarity and retrieve the associated values stored in the EM. The hidden states (ℎ_*t*_) generated from the RNN serve as values (𝑣_*t*_ = ℎ_*t*_), representing memory contents and are encoded in an EM buffer at every time step. Linear transformations are denoted with 𝑾_𝒌_ and 𝑾_𝒒_ ∈ ℝ^2𝑑×𝑑^, following Gershman et al.^30^.

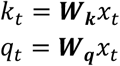

The key and value at every time step were encoded in the EM, with the maximum number of memory (𝑀) set as the number of time steps in test data.

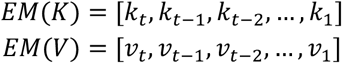

The query surveys all keys stored in EM with the following equation. The equation assigns high attention weights to keys that are similar in pattern with the query, with the softmax function making the attention weights sum to 1.

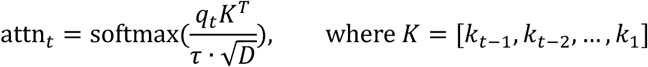

𝜏 was set to 0.1 to enforce sparsity in memory selection—meaning, higher attention weights were given to a small number of past scenes while most of the memories were given near zero weights. Because the dimension of the hidden state (𝐷) was 100, the attention calculation essentially equaled to: attn_*t*_ = softmax(𝑞𝐾^𝑇^). Attention is calculated from the match between key and query, but the model retrieves values stored in the EM that are associated with the attended keys. The retrieved memory at time *t* is denoted as 𝑚_*t*_ ∈ ℝ^2𝑑^, which is the summation of past values, weighted by the model-learned self-attention.

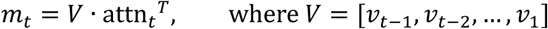

The outcomes of the illustrated steps are ℎ_*t*_ representing the current scene and 𝑚_*t*_ representing the retrieved and integrated past memories. We integrated the two representations by a weighted average of the two vectors: ℎ_*t*_ × 𝛼 + 𝑚_*t*_ × (1 − 𝛼). Here, 𝛼 was set to 0.5, which means the two vectors were simply averaged in memory integration. Given this integrated vector, model parameters 𝑾_𝒚_ ∈ ℝ^𝑑×2𝑑^ and 𝒃_𝒚_ ∈ ℝ^𝑑^ were fitted to predict the embedding of the next time step 𝑥_*t*+1_.

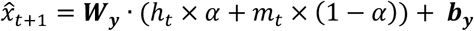

The model parameters were initialized as follows. Weight parameters 𝑾_𝒙_, 𝑾_𝒉_, and 𝑾_𝒚_ were initialized using orthogonal initialization, and bias parameters 𝒃_𝒙_, 𝒃_𝒉_, and 𝒃_𝒚_ were initialized with a constant value of 0.1. For the input-to-key and input-to-query transformation matrices, 𝑾_𝒌_ and 𝑾_𝒒_, we used trainable parameters initialized with Gaussian noise scaled by a factor of 0.1.

The model was trained using an Adam optimizer (torch.optim.Adam) with a learning rate of 0.001. The training objective was to minimize the mean squared error between the predicted and observed vectors (nn.MSELoss).

The model was trained 50 times, with each iteration comprising weight optimizations with 17 episodes’ embedding time series. In total, this amounted to 850 weight updates (17 episodes × 50 iterations). 17 episodes were presented at a random order at every iteration. We repeated this entire training and testing steps 20 times with different random seeds.

Model parameters were fixed at every round of training iteration to be applied to test data. Test was conducted on three different scrambling orders of episode 1. The results for these three test runs were averaged, upon unscrambling the events into their original sequence.

Performance of the EM-GRU model was compared to five other comparison models. First, in *shuffled memory* model, everything was the same as the original EM-GRU except that the attention weights (attn_*t*_) calculated at every time step were randomly shuffled when they were applied to the stored values in the EM. This means randomly selected memories were retrieved as 𝑚_*t*_. Second, in *no key* model, the query directly searched through the stored hidden states, thus the self-attention calculation changed as follows.

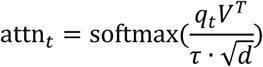

Third, in *no key-query* model, neither key nor query existed, thus the self-attention calculation was based on the following equation.

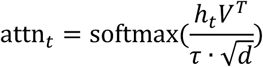

Here, if the 𝑉 were to include the nearby time steps (e.g., ℎ_*t*−1_, ℎ_*t*−2_), their pattern similarity with ℎ_*t*_ was bound to be high because ℎ_*t*_ is dependent on the recurrence from the nearby past time steps. Therefore, we only included memories that were 10 time steps apart so that the model could focus on retrieving memories from distant past.

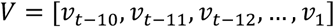

Fourth, the *no EM (GRU)* model was implemented as a traditional GRU RNN that does not have the EM buffer. From the input 𝑥_*t*_, the model estimates the hidden representation ℎ_*t*_ with recurrence, and from this hidden state, directly predicts the next input 𝑥_*t*+1_.

Finally, to implement the *no WM* model, the GRU was lesioned and replaced with a multilayer perceptron, such that ℎ_*t*_ = 𝑾_𝒙_𝑥_*t*_ + 𝒃_𝒙_ (𝑾_𝒙_ ∈ ℝ^2𝑑×𝑑^ and 𝒃_𝒙_ ∈ ℝ^2𝑑^). The EM buffer was identical to that used in the EM-GRU model.

### Human behavioral and fMRI data

Detailed descriptions of the experiments and data are provided in Song et al.^12^. Each fMRI session consisted of 10 runs. During each run, 36 participants watched several events while pressing an “aha” button whenever they understood something new about the story or characters. Afterward, they were shown screenshots of when they pressed the button and participants verbally explained why they pressed in those moments.

To create a memory retrieval matrix, we analyzed people’s verbal responses explaining their insights. 41.15 ± 16.64 % of verbal responses included mentions of past events. For those cases, we noted an event index of the mentioned past event, an event index of the current moment when they had an “aha,” and recorded +1 for that event pair. This resulted in one memory retrieval matrix per person. Upon unscrambling the events to their original order, we averaged all 32 participants’ data (4 participants’ data were missing due to errors in data collection) to create one memory retrieval matrix that was used in this study. Memory retrieval at events 46 to 48 were omitted during coding, so they were discarded when calculating correlations. To estimate the noise ceiling for the model and human participants’ memory retrieval similarity, we adopted a split-half reliability approach. We randomly split 32 participants into two groups, computed the average retrieval matrix within each group, and calculated the correlation between the two group-averaged matrices. We repeated this procedure 10,000 times with different random splits to estimate one mean *rho* value, which served as the noise ceiling estimate.

The causal relationship matrix was not collected from this fMRI sample but was rated by three authors of the Song et al.^12^ study, one of whom is the first author of this study. Each event pair was assigned a causal relationship score from 0 to 4. Because the scores were highly reliable across the raters, the three ratings were averaged to create one causal relationship matrix that was used in this study.

### Event-by-event representation pattern similarity

The representation patterns of hidden states estimated across time (100 dimensions × time) were summarized to a matrix sized (100 dimensions × 48 events) by averaging patterns within each event. Representation pattern similarity analyses were conducted by taking the Pearson’s correlation between events (48 events × 48 events), which were Fisher’s z-transformed. Event orders were unscrambled to follow the original order of the episode, and the RSMs of the three scrambled order groups were averaged.

In correlation analyses related to **Figures 4** and **5**, we controlled for the input pattern similarity matrix using partial correlation. This was because input similarity matrix was significantly positively correlated with both human memory retrieval matrix (𝜌 = 0.472, *p* = 5e-56) and causal relationship matrix (𝜌 = 0.542, *p* = 3e-87). This approach regresses out the semantic and perceptual feature similarities in event pairs that are explicitly given by the input CLIP embeddings. To calculate the noise ceiling of the partial correlation, we controlled for the input pattern similarity in the split-half reliability analysis.

Following the same procedure used for the model, we created brain RSMs from the voxel activity pattern time series. The BOLD activity time series were extracted from all voxels in each of the 200 cortical^47^ and 32 subcortical^48^ parcels as participants watched 48 events in scrambled orders. The BOLD time series were averaged across time for 48 events respectively, resulting in a (number of voxels × 48 events) representation pattern matrix per region. We created 33 participants’ RSMs (3 participants’ data were omitted due to head motion) for each of the 232 brain regions, and unscrambled the event orders. To estimate the similarity between the brain RSM and the model RSM, we calculated the correlations between each participant’s brain RSM and the model RSM while controlling for input similarity. The resulting correlation values were Fisher *z*-transformed and averaged across 33 participants. Statistical significance was assessed by comparing this value to a null distribution generated by randomly shuffling the brain RSMs (1,000 iterations). Multiple comparisons were corrected across 232 brain regions. To compare the EM-GRU and GRU models in terms of their similarity to the brain RSMs, we conducted paired *t*-tests using the Fisher’s *z*-transformed correlation values between the 33 brain RSMs and the model RSMs derived from either the EM-GRU or GRU.

## Supplementary Information

### Supplementary Figures

**Supplementary Figure S1.**
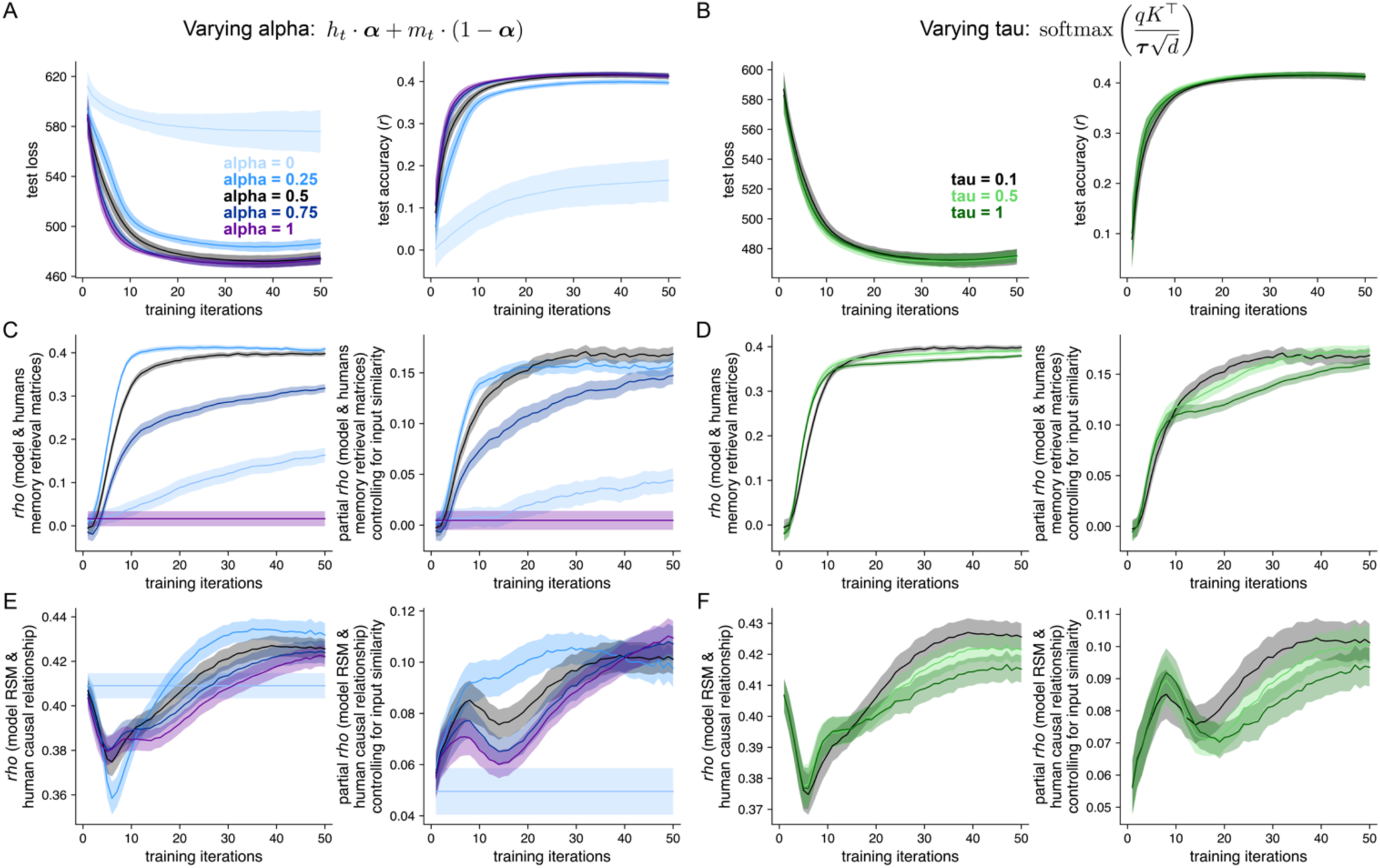
EM-GRU results across varying hyperparameters 𝛼 and 𝜏. All other aspects of the model were identical to the original EM-GRU model. Black lines correspond to the EM-GRU results reported in the manuscript (𝛼 = 0.5 and 𝜏 = 0.1). **(A-B)** Performance on the next-scene prediction task, complementing Figure 3. **(A)** Model performance improves with increasing 𝛼. This indicates that the next-scene prediction is more suaccessful when predictions rely more on ongoing scenes rather than retrieved memories. **(B)** Model performance is comparable across different choices of 𝜏. **(C-D)** Comparison between the humans and model’s memory retrieval, complementing Figure 4. **(C)** The model retrieves memories more like humans on the order of 𝛼 = 0.5, 0.25, 0.75, 0, and 1. This indicates the importance of integrating ongoing event representations with retrieved memories. **(D)** Models with varying 𝜏 retrieve memories like humans to a similar extent. **(E-F)** Comparison between the model’s representation similarity matrices (RSMs) and the human causal relationships, complementing Figure 5. **(E)** Models with varying 𝛼 represent events based on causal event structure to a similar extent, near training iteration = 35. **(F)** The model represents events more based on causal event structure on the order of 𝜏 = 0.1, 0.5, and 1. This indicates that attending to selective memories by giving sparse attention weights is more human-like than broadly attending to a large number of memories.

**Supplementary Figure S2.**
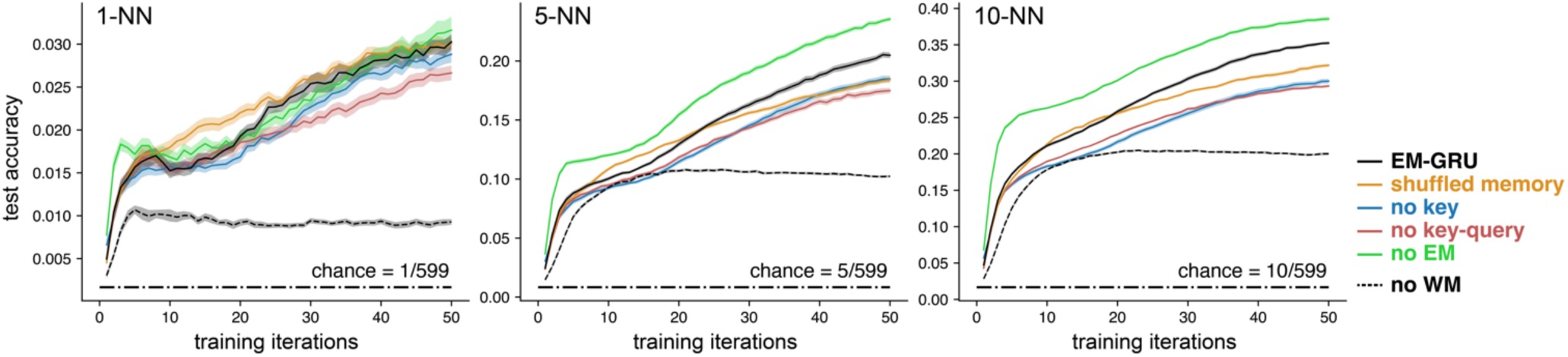
Validation of model performance using a nearest-neighbor (NN) accuracy metric. In the main manuscript, model performance was quantified as the average Pearson’s correlation between the predicted and observed CLIP embedding vectors of the next time steps. To assess robustness, we additionally evaluated performance using a NN accuracy metric at test. This metric computes the percentage of time steps in which the predicted embedding for the next time step falls within the top-*N* highest Pearson’s correlations among all 599 possible scenes. We report results for top-1 (*left*), top-5 (*middle*), and top-10 (*right*) NN accuracy. Conditions match those shown in Figure 3. Dot-dashed lines at the bottom indicate chance performance: *N* out of 599 possible scenes.

**Supplementary Figure S3.**
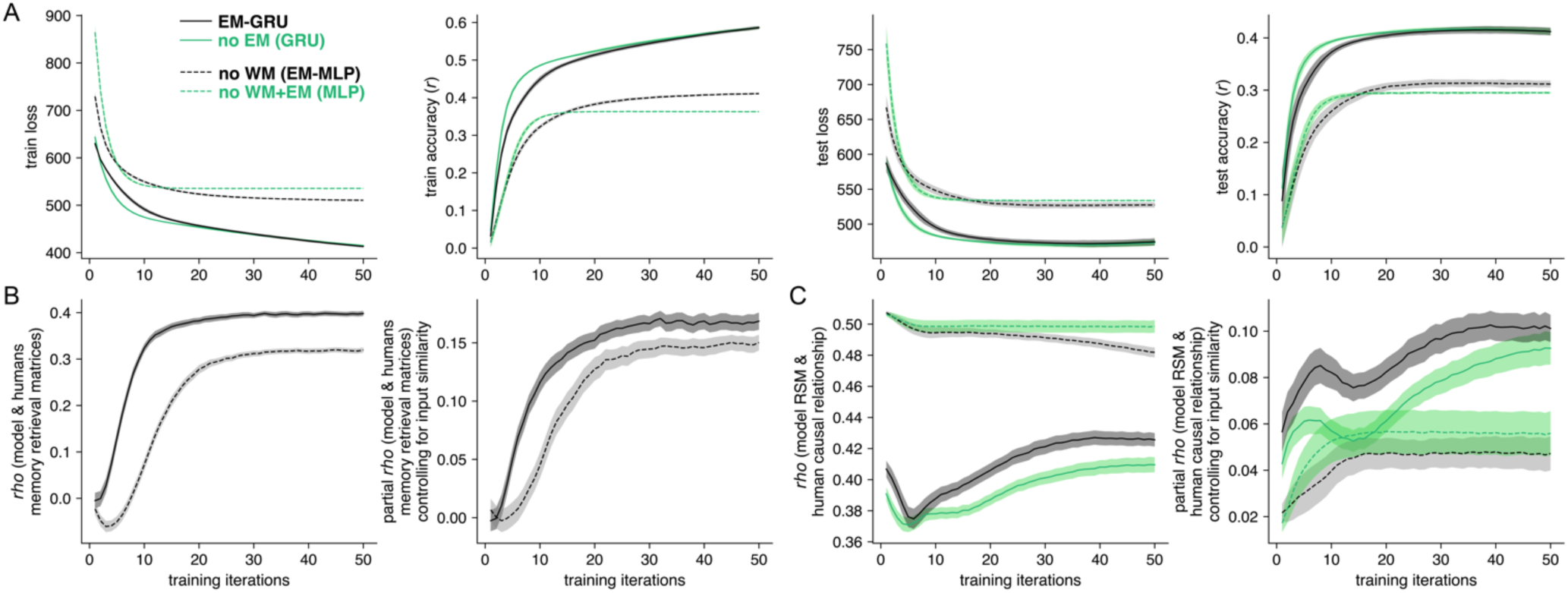
The role of working memory (WM) supported by recurrence in the gated recurrent unit (GRU) recurrent neural network (RNN). In the manuscript, model manipulations targeted the episodic memory (EM) buffer, because our goal was to understand how people organize and retrieve long-term episodic memories. In this figure, we instead manipulated the GRU RNN to assess the role of working memory within the same movie-watching context. Specifically, we replaced the GRU with a multilayer perceptron (MLP) while matching the size of the hidden layer. With respect to the EM-GRU (black solid line) and GRU (*no EM*; green solid line) models reported in the main manuscript, we present the performance of the EM-MLP (*no WM*; black dashed line) and MLP (*no WM+EM*; green dashed line). **(A)** Performance on the next-scene prediction task, complementing Figure 3. Replacing the GRU with an MLP substantially reduced performance on the next-scene prediction task. When considering that the models of EM variants did not vary in explicit task performance, this result suggests that the recurrence in the GRU is mainly responsible for the next-scene prediction task. **(B)** Comparison between the humans and model’s memory retrieval, complementing Figure 4. The EM-GRU’s memory retrieval was more similar to humans’ than the *no WM* model’s. **(C)** Comparison between the GRU and MLP models’ representation similarity matrices (RSMs) and the human causal relationships, complementing Figure 5. When considering raw correlations, the MLP-based models’ representation pattern similarity (RSM) more closely resembled the events’ causal relationships. However, after controlling for the input pattern similarity, the GRU-based models represented causally related events with more similar representations. Together, the results suggest that recurrence in the GRU contributes to both explicit task performance and emergent memory representation and retrieval.

**Supplementary Figure S4.**
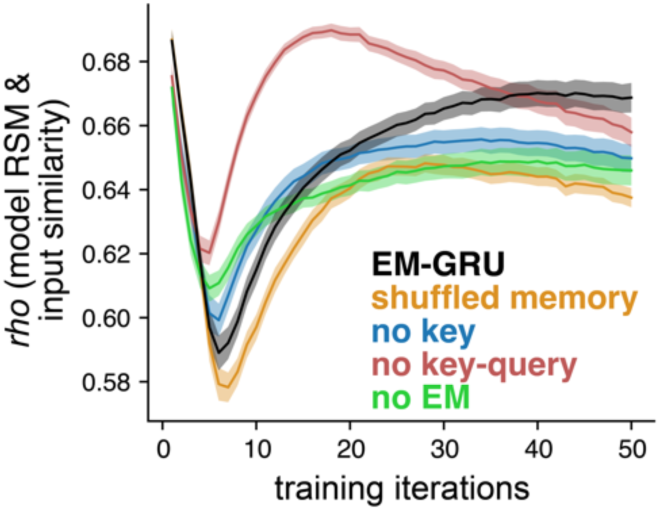
Correlation between the model’s hidden state representation pattern similarity (RSM) and input pattern similarity. The figure complements Figure 5B-C.

**Supplementary Figure S5.**
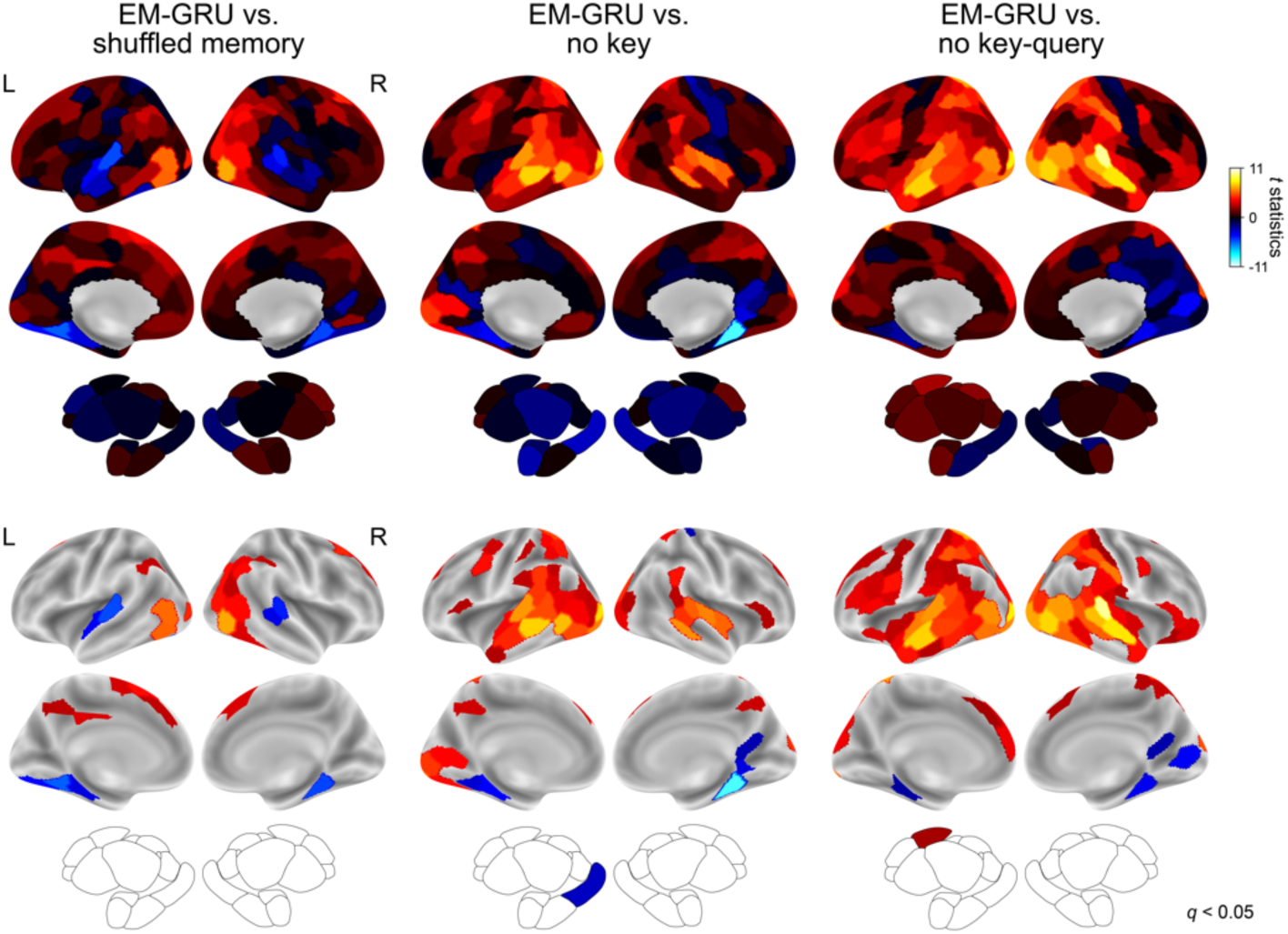
Comparisons between the EM-GRU and control models in how much model representation similarity matrices (RSMs) resemble brain region RSMs. The figure complements Figure 6B, which reports paired *t*-test results comparing the correspondence between the brain RSMs and EM-GRU vs. GRU model RSMs. Here, we show results for other control models: the *shuffled memory*, *no key*, and *no key-query* models. In nearly all cases, EM-GRU representations show higher correspondence with the human brain representations (indicated in red-yellow). The top panels display raw *t* statistics and the bottom panels show the same results after thresholding at FDR-corrected *p* < .05.

**Supplementary Figure S6.**
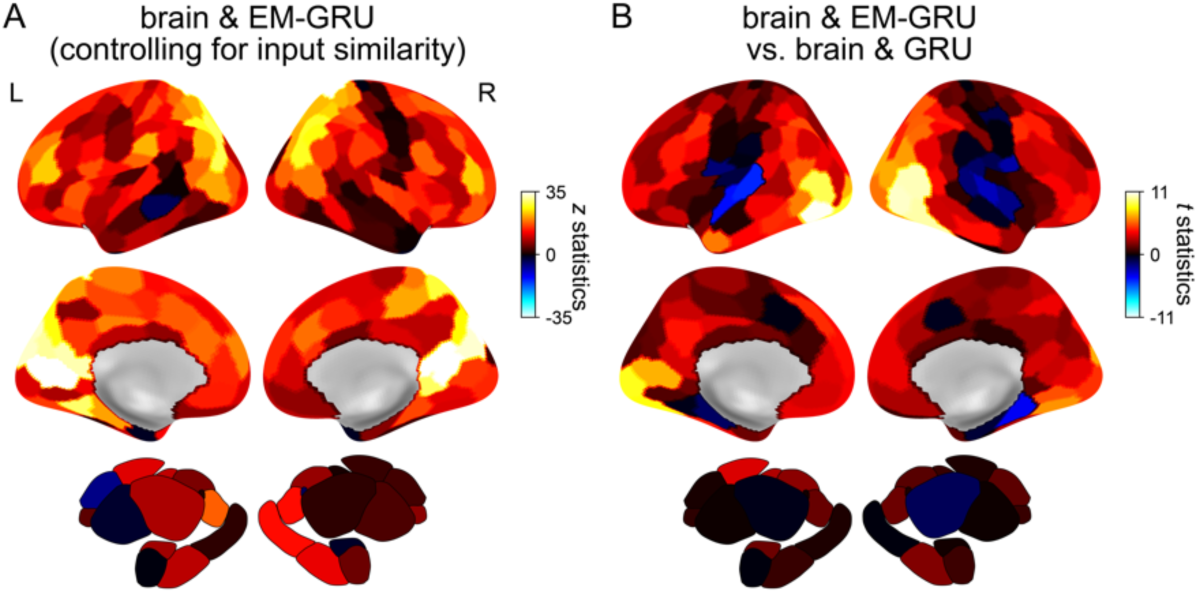
Unthresholded visualization of Figure 6.

**Supplementary Figure S7.**
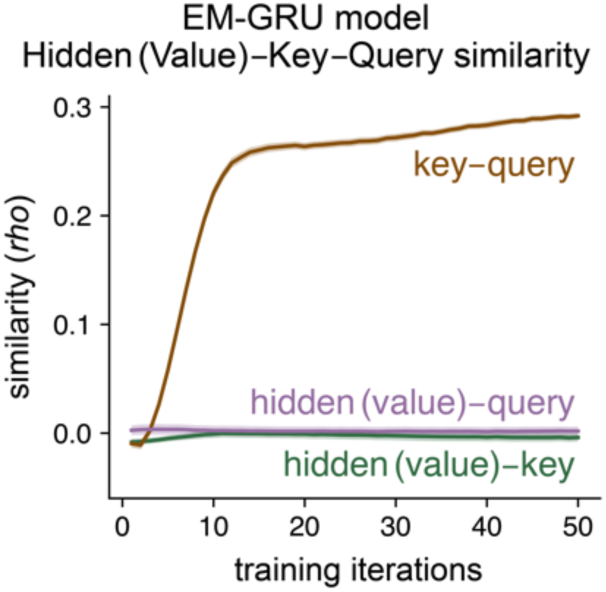
Pairwise similarity between the hidden state (value), key, and query of the EM-GRU. Lines indicate the mean and error bars indicate the standard error of the mean across 20 repetitions using different random seeds.

### Supplementary Texts

**Supplementary Text S1.**
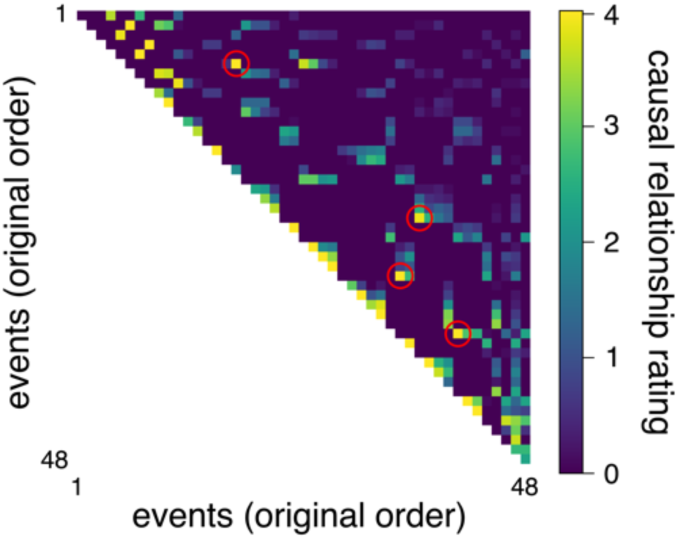
Examples of causally related events The following descriptions correspond to event pairs that were consistently rated as highly causally related by all three raters. The descriptions match the red-circled event pairs shown in the figure, from top to bottom.

#### Pair 1

[Event A] Randall is in his office. He opens an email titled “GOOD NEWS”. The message says they have found a guy named Willian Hill, including his age and home address. On his laptop screen is a photo of an elderly Black man. Randall looks emotional.

[Event B] Randall is at his daughters’ soccer game with his wife. As they watch the daughters play, Randall tells her that he found his father by hiring a guy. His wife asks what he’s going to do, and Randall replies he doesn’t know. He says his birth mother was a crack addict who died during childbirth, and his birth father left him at a fire station. When his wife asks, “Then why did you find him?” Randall replies, “I don’t know” and looks emotional. Their daughter scores a goal and they both get excited.

#### Pair 2

[Event A] Kate is pouring coffee in her paper cup when Toby comes and says hi. They chat about what had happened, chuckle, and introduce themselves. Toby blurts out “So, you wanna be fat friends?” and Kate laughs and says sure. They are flirting. Kate says she’s determined to lose the weight, but Toby says he probably is not. Kate says she can’t fall for a fat person right now. Toby responds, “Well… I guess I’ll lose the weight then” and leaves. Kate blushes.

[Event B] Kate and Toby are on a date in a restaurant. They are talking about silly things and are laughing together and having fun. The server approaches the table and asks whether the couple wants desserts. Toby says yes, Kate says no, and they go back and forth. Toby says “I am utterly fascinated by dessert. Dessert is my life’s work.” When Kate finally gives up, Toby says “We will just have the check, please.”

#### Pair 3

[Event A] Randall is having a serious talk with William in a small dark room. William talks about the fire station and how his life is a punishment already, and Randall listens. Randall looks angry and says he’s not going to forgive William. He says he just wants to say, “Screw you” and storm out of there. William says, “Go ahead”, so Randall shouts, “Screw you” and storms out of the room. Soon after, Randall comes back. Randall asks, “You want to meet your grandchildren?” William looks hopeful and replies, “I’ll get my coat.” [Event B] Randall drives to a big house. When they enter, Randall is greeted by his two daughters and his wife. The wife looks startled to see William. Randall introduces William to everyone. William shakes his hands with Randall’s wife. The daughters say that William has a hole in his pants, and they start talking. William makes the daughters laugh.

#### Pair 4

[Event A] Rebecca is in labor. She is sweating and screaming in the hospital room and the doctor and nurses are asking Rebecca to push. The first baby boy is out, but Rebecca says she can’t breathe. Rebecca is put on an oxygen mask, and the doctor says he’s going to take it from now. The doctor orders Jack to get out of the room, and Jack resists, saying that he needs to be with his wife. It’s an emergent situation, Rebecca is unconscious, and Jack is kicked out of the room.

[Event B] Jack is waiting in the waiting area. The doctor, dressed in a surgery gown, walks toward Jack. The doctor says Rebecca is fine, they had another baby girl, but they lost the third child. Jack is in shock and can’t process the news. The doctor explains the situation again. Jack is trembling and says he needs to be with his wife. The doctor gently stops him and asks him to sit down.

## Notes

### Competing Interest Statement

The authors have declared no competing interest.

### Summary of Updates

The manuscript has been revised overall.

